# Cdh2 coordinates Myosin-II dependent internalisation of the zebrafish neural plate

**DOI:** 10.1101/424705

**Authors:** Claudio Araya, Hanna-Maria Häkkinen, Luis Carcamo, Mauricio Cerda, Thierry Savy, Christopher Rookyard, Nadine Peyriéras, Jonathan D Clarke

## Abstract

Tissue internalisation is a key morphogenetic mechanism by which embryonic tissues generate complex internal organs and a number of studies of epithelia have outlined a general view of tissue internalisation. Here we have used quantitative live imaging and mutant analysis to determine whether similar mechanisms are responsible for internalisation in a tissue that apparently does not have a typical epithelial organisation – the zebrafish neural plate. We found that although zebrafish embryos begin neurulation without a conventional epithelium, medially located neural plate cells adopt strategies typical of epithelia in order to constrict their dorsal surface membrane during cell internalisation. Furthermore, we show that Myosin-II activity is a significant driver of this transient cell remodeling which also depends on Cdh2 (N-cadherin). Abrogation of Cdh2 results in defective Myosin-II distribution, mislocalised internalisation events and defective neural plate morphogenesis. Our work suggests Cdh2 coordinates Myosin-II dependent internalisation of the zebrafish neural plate.

## Introduction

The internalisation of superficial sheets of cells is a widely used developmental strategy to generate complex three-dimensional structures with well-defined shape and size. Through this mechanism, animal tissues form a number of internal organs including the vertebrate central nervous system^1,2^. In recent years, live imaging studies and mutant analysis has begun to define the cellular, molecular, and biomechanical mechanisms responsible for tissue internalisation in a growing number of tractable model systems^3,4^. Collectively, these studies have identified that tissue internalisation is thought to occur by an interplay of cell shape changes, cell adhesion remodelling and subcellular cytoskeletal protein dynamics, including the cortical Actomyosin motor complex^2,4^. In recent years, Drosophila ventral midline furrow formation during gastrulation has been an instrumental model system to dissect the cellular and molecular mechanisms underlying *in vivo* tissue internalisation^5,6,7^. Live imaging analysis in gastrulating flies have indicated that tissue internalisation is achieved by a coordinated activity of medial cells which show progressive and irreversible cell surface constriction while keeping a more or less constant cell volume^6,8^. Furthermore, recent studies have demonstrated that this cell behaviour is powered by cortical Myosin-II network^7^, and that the cell-cell adhesion molecules including E-Cadherin are critical to efficiently transmit and coordinate tension across the internalising tissue^9^. Thus apical constriction has been identified as a dominant and instrumental cell behaviour for surface tissue internalisation in epithelia.

Neurulation in zebrafish is a complex morphogenetic event that first transforms the neural plate into a neural keel and then a neural rod before lumen formation generates the neural tube structure. The details of this process are incompletely understood but initially involve two components, one is convergence of neural plate cells towards the midline and the second is an internalisation of cells at or close to the midline^10,11^. The efficiency of convergence depends on Planar Cell Polarity signaling^12,13,14^ and requires extracellular matrix and adjacent mesoderm for coordination^15,16^. Internalisation is less well understood but is a key step that deepens the most medial zone of the neural plate to generate the solid neural keel. While the most medial cells of the plate are internalising the more lateral cells are still converging to the midline to take the place of the internalised cells. In this respect the tissue movement appears somewhat like a conveyor belt, narrowing the neural plate as it deepens medially. The cell behaviours that underlie this tissue movement are not fully understood, however they are not simple and likely involve cell shape changes, cell orientation changes and cell intercalations. During this period of internalisation the cells of the neural plate and keel are not organised as a columnar neuroepithelium as found in other vertebrates. The pseudostratified epithelial organisation does not arise in teleosts until late neural rod stage, coincident with lumen formation^12,13,14,15,16,17,18,22^. This is in contrast to amniote and amphibian neural plates that have a clear epithelial organisation and use apical constriction to fold the epithelium and internalise the neuroectoderm during neurulation^19,20^. This poses the question of what cell behaviours drive internalisation in the fish neural plate. So far the best clue to this is the dependence of this process on the cell adhesion protein Cdh2 (previously called N-cadherin). Embryos mutant for Cdh2 fail to complete convergence and internalisation of the neural plate, with the phenotype particularly strong in the hindbrain region^21,22^. A reduction in protrusive behavior of neural plate cells has been suggested to contribute to this phenotype^22^ but Cdh2-dependent convergence and internalisation remains incompletely understood.

Here we have applied quantitative live imaging and genetic analysis to understand tissue internalisation in the hindbrain region of the zebrafish neural plate. We show that while the organisation and movements of the teleost neural plate are distinct from neural plate in other vertebrates, cell internalisation at the dorsal midline is achieved by adopting similar cellular strategies. This includes deployment of Cdh2 and Myosin-II to effect constriction of the dorsal cell surfaces to generate inward traction. Furthermore, we show this medial neural plate behaviour depends on Cdh2 function and superficial non-muscle Myosin-II activity at the internalisation zone. While Myosin-II inhibition blocks cell surface constriction and cell internalisation, depletion of Cdh2 leads to mislocalised Myosin-II distribution and random cell internalisation events along the dorsal surface. Together, these results suggest the zebrafish neural plate deploys strategies of cell surface constriction similar to conventional epithelia to effect internalisation. Overall, our observations suggest Cdh2 coordinates Myosin-II dependent internalisation of the zebrafish neural plate.

## Results

### Neural plate internalisation occurs through reorientation and elongation of neural plate cells

In the prospective hindbrain region, the zebrafish neural plate is a multi-layered tissue of 3-6 cell deep at 10 hours post fertilisation (hpf)^12,15^ (Fig. 1a timepoint 0 min). To study changes in cell morphology we first labelled cells with plasma membrane constructs and made confocal time-lapse movies in the transverse plane. At the 10 hpf stage, most cells are largely round-shape in transverse section and progressively elongate in the transverse plane as they converge to the midline and internalise to form the neural keel (Fig. 1a,b,c and Movie 1). To quantify these cell shape changes we assigned cells to lateral zones (between 20-60 μm from the midline) and medial zones (within 20 μm of the midline) of the neural plate, and then measured the long axis of cells and the angle they subtend to the parasagittal plane of the embryo (Fig. 1c,d). Up to 11 hpf mean cell length increases at the same rate in both lateral and medial zones, after 11 hpf cells in the medial zone continue to elongate while the mean length of more lateral cells is more stable (Fig. 1d). The orientation of the long axis of both medial and lateral cells leans in towards the midline at an angle of approximately 50° up to 11 hpf, after that time medial cells reduce this angle to approximately 30° while lateral cells increase their angle to approximately 80°. Between 11 and 13 hpf the depth of the neural plate increases in the medial zone by approximately 300% (from 42μm to 138μm on average, Fig. 1a,b, and Movie 1, n_embryos_ = 4) as medial cells internalise and the plate begins to transform into the neural keel. To analyse the overall coordination of hindbrain neural plate cells during internalisation we made movies from 9-11 hpf of cells labelled with the nuclei marker H2B-GFP (Movie 2). We then tracked nuclei movements using 3D+time tracking analysis on the Bioemergence platform + Mov-IT pipeline^23^ (Fig. 1e-g, n_embryos_ = 3). We first assessed and quantified nuclei movements from the dorsal view (Movie 2). Up to 11 hpf nuclei are seen to move both medially and anteriorly (Fig.1e). From 11 hpf nuclei move more directly medially and the anterior component to their movement is largely lost (Fig. 1e’). We next tracked nuclei in the transverse plane and between 9 and 11 hpf most nuclei move medially with relatively straight trajectories towards the midline (Fig. 1f,g). Between 11 and 12 hpf nuclei close the midline change their direction and begin to move deeper into the embryo. Between 12 and 13 hpf most nuclei within 40 μm of the midline are directed down into the embryo. While medial nuclei move deep into the embryo, more lateral nuclei are still moving medially as they converge towards the midline. An analysis of speed shows lateral cells accelerate towards the midline between 10 and 11.5 hpf, while medial cells initially move quicker than lateral cells but after 11 hpf move more slowly (Fig. 1h) (n_embryos_ = 3, n_average of medial cells analysed_ = 90, n_average of lateral cells analysed_ = 80 in each transverse view assay, P<0.0001). In conclusion, convergence towards the midline is accompanied by an elongation and reorientation of cells, and internalisation involves a distinctive reorientation of nuclei movements in the medial zone of the neural plate.

**Figure 1.**
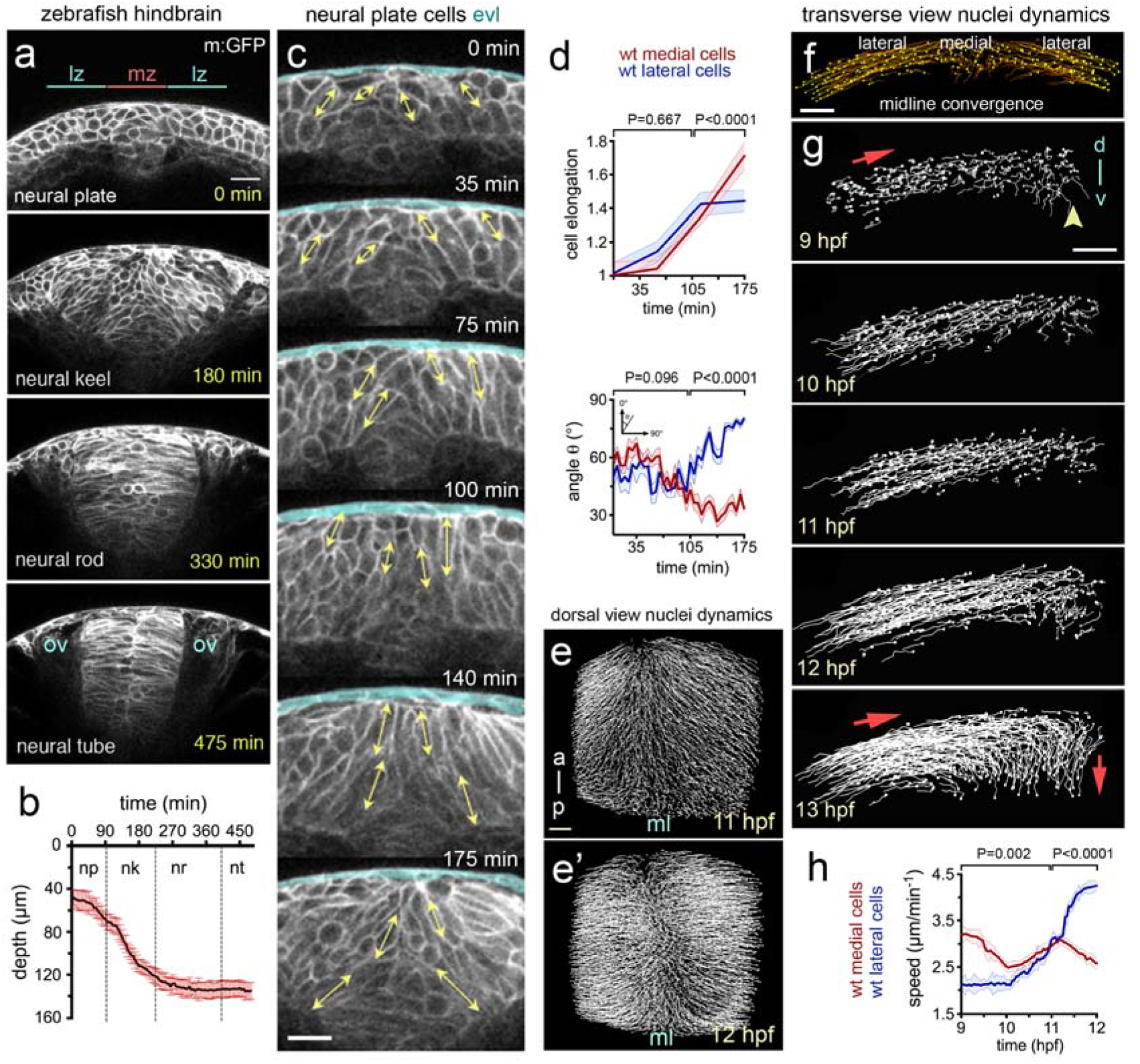
Cell and tissue dynamics during zebrafish neural plate internalisation. (**a**) Four transverse view confocal images depicting the neural plate, keel, rod and tube stages of wild-type zebrafish neurulation at hindbrain levels. Cells labelled with membrane-GFP (m:GFP). Scale bar is 20μm and OV indicates otic vesicle as hindbrain landmark. The medial zone (mz) and lateral zone (lz) of the neural plate are indicated in (a) and each is 20 μm wide. (**b**) Time course showing depth of the neural anlage at the midline. Internalisation deepens the neural anlage by approximately 300% in the plate to keel transition (initial thickness ~ 42μm on average vs final thickness ~138μm on average, n_embryos_ = 4). np, neural plate; nk, neural keel; nr, neural rod; and nt, neural tube. (**c**) Frames from transverse view time-lapse to illustrate cell profile changes during plate to keel transition. Double headed arrows illustrate examples of measurements of the long axis of cells used to quantify cell elongation. EVL (enveloping cell layer) is highlighted in pale blue. (**d**) Quantification of cell elongation and cell angle in medial and lateral zones during plate to keel transition. (**e and e’**) Projected views from dorsal surface of nuclei tracks at 11 hpf and 12 hpf. Nuclei tracked from movies of cells labelled with H2B-GFP (**f**) Transverse view image of nuclei tracked during convergence to midline (10.5 hpf). Scale bar, 20μm. (**g**) Transverse views of nuclei tracks of neural plate dynamics during convergence and internalisation (9-13 hpf, for simplicity only half of the neural plate is illustrated). Red arrows indicate tissue movement, and arrowhead indicates midline position. D, indicates dorsal, while v, indicates ventral. (**h**) Quantification of nuclei speeds for medial and lateral cells during convergence and internalisation (n_embryos_ = 3, n_average of medial cells analysed_ = 90, n_average of lateral cells analysed_ = 80, 5 z-slice analysed in each embryo).

### Internalisation of neural plate cells is achieved by progressive cell surface reduction

To better understand neural plate cell internalisation, we imaged membrane labelled cells from dorsal to obtain an *en face* surface view of hindbrain neural plate (Fig. 2a). Using z-stacks with 0.25μm z-intervals, we could eliminate dorsal enveloping layer cells (EVL) from above the neural cells to reveal the cell dynamics at the dorsal surface of neural plate. At 11-12 hpf, following individual medial cells showed that these cells undergo progressive dorsal surface reduction and internalisation and eventually become lost from the imaging plane (Fig. 2b). Most individual lateral cells largely do not reduce their dorsal profile over this time (Fig. 2b). This was confirmed by quantifying the dorsal cell profiles within a medial and a lateral region of interest over a 60 minute time period (Fig. 2c, Movie 3 and Movie 4). In general the dorsal profiles in the medial zone are smaller than the profiles in the lateral zone, but there is some heterogeneity in both (small profiles are darker grey than the larger profiles in Fig. 2c). Over this time period the mean dorsal surface profile of medial cells decreases by approximately 50%, while the mean of cells in the lateral zone remains constant (Fig. 2d). Contraction of the superficial dorsal surface of cells thus occurs largely in the medial zone of the neural plate and largely occurs on a cell by cell basis as we find small profiles surrounded by large profiles in this zone. To quantify cell surface constriction we measured both cell aspect ratio (independent of cell orientation), and cell surface alignment relative to the embryo axis (Fig. 2e, f, n_embryos_ = 4, ∼75 medial and ~70 lateral cells in each assay). These data show the dorsal surface of cells tends to constrict anisotropically (more in the mediolateral dimension than their anteroposterior dimension) and the long axis of this surface of the cells is weakly aligned to the anteroposterior axis of the embryo. Taken together, these analyses demonstrate that neural plate internalisation is preceded by progressive anisotropic reduction of the dorsal surface of cells in the medial zone.

**Figure 2.**
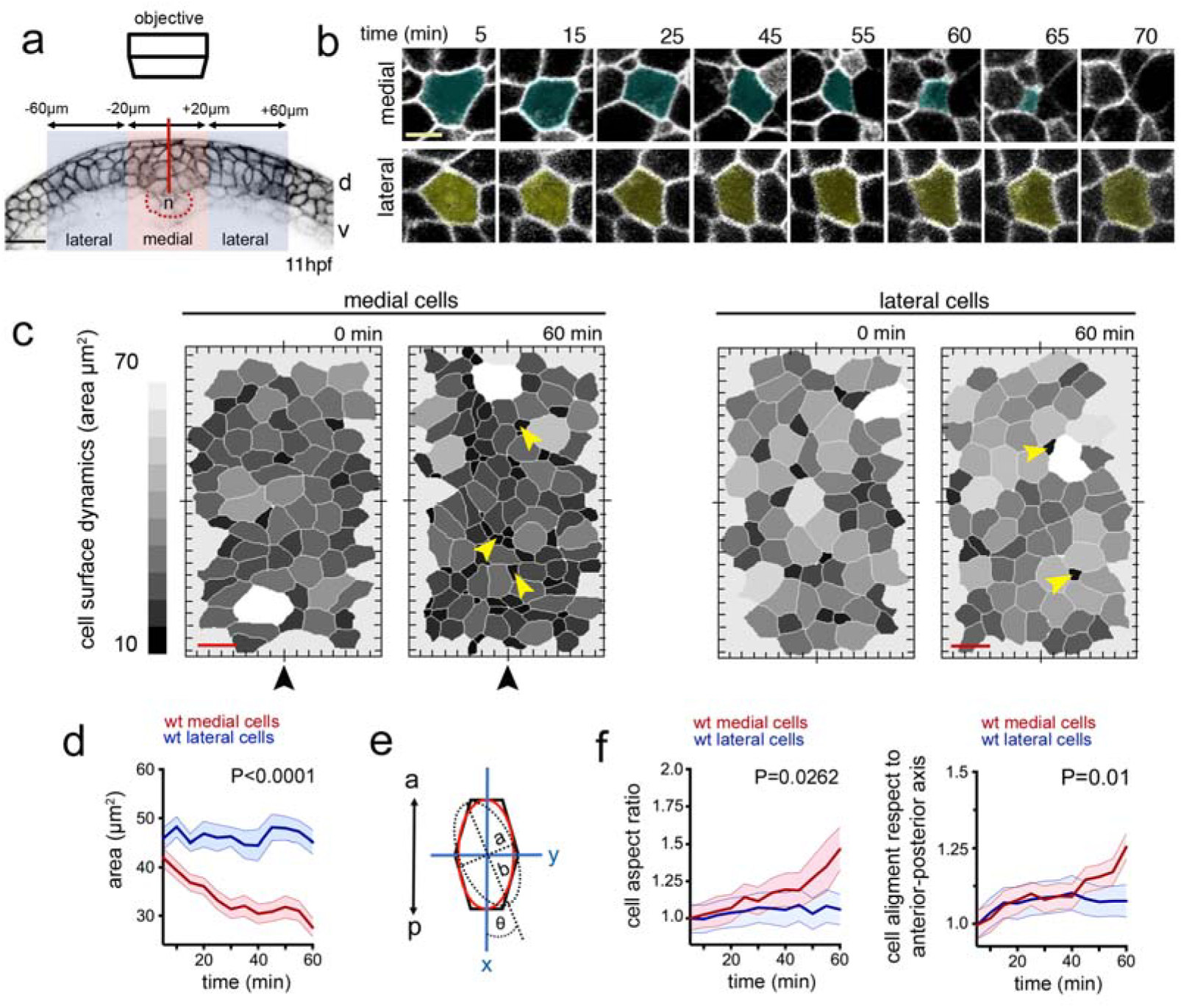
Dorsal cell surface dynamics during internalisation. (**a**) Schematic of the imaging approach to visualise dorsal cell surface profile during convergence and internalisation. Inverted LUT image depicts cell membrane outlines. D is dorsal, v is ventral, and n is notochord. Scale bar is 20μm. (**b**) Tangential z-slices 5-7 μm below EVL were selected for automated segmentation of dorsal surface of superficial neural plate cells. Top panels: image of dorsal surface of medial cell over time (pseudocoloured in cyan). Bottom panels: images of dorsal surface of lateral cell over time (pseudocoloured in yellow). Scale bar, 5μm and time is in minutes. (**c**) Dorsal surface profiles of medial and lateral populations of cells one hour apart. For simplicity only half of the neural plate is shown. Colour code indicates relative dorsal surface profile area in μm ^2^, darker greys are smaller than lighter greys. Black arrowhead indicates midline position. Scale bar, 10μm. (**d**) Mean dorsal surface area of medial (red) and lateral (blue) cells over time (n_embryos_ = 4 wt analysed, P<0.0001). (**e**) Schematic representation of a cell’s dorsal surface area to illustrate cell aspect ratio and alignment of this surface area to the anteroposterior axis of the embryo. (**f**) Quantification of surface aspect ratio and alignment to embryonic axis. In all graphs error bars indicate SEM.

### Myosin-II is required for zebrafish neural plate internalisation

In other systems, apical surface constriction leading to internalisation is thought to be powered by apical actomyosin networks^3,4,7^ To assess whether myosin activity is likely to drive contraction of the dorsal surface of medial neural plate cells we first used immunohistochemistry to analyse the localisation of phospho-Myosin light chain. Consistent with a role for myosin in surface constriction we detected high levels of phosho-Myosin along the dorsal surface of superficial neural plate cells immediately underneath the large non-neural EVL cells (Fig. 3a-d). Distinctive phospho-Myosin puncta could also be seen deeper in the neural plate scattered along the Phalloidin stained cell cortices (Fig. 3e-g’’). We next checked whether we could monitor actomyosin localisation in the neural plate in living embryos to determine its dynamic distribution during neural plate morphogenesis. The transgenic embryos (Tg(*actb1:myl12.1-*GFP), hereafter called Myosin-II:GFP) and (Tg(*actb1*:GFP-*utr*CH), hereafter called Actin:GFP)^24,25^ faithfully recapitulated the distribution of phospho-Myosin and F-actin in neural plate cells revealed by antibody and Phalloidin staining respectively (Fig. 3h-j’’).

**Figure 3.**
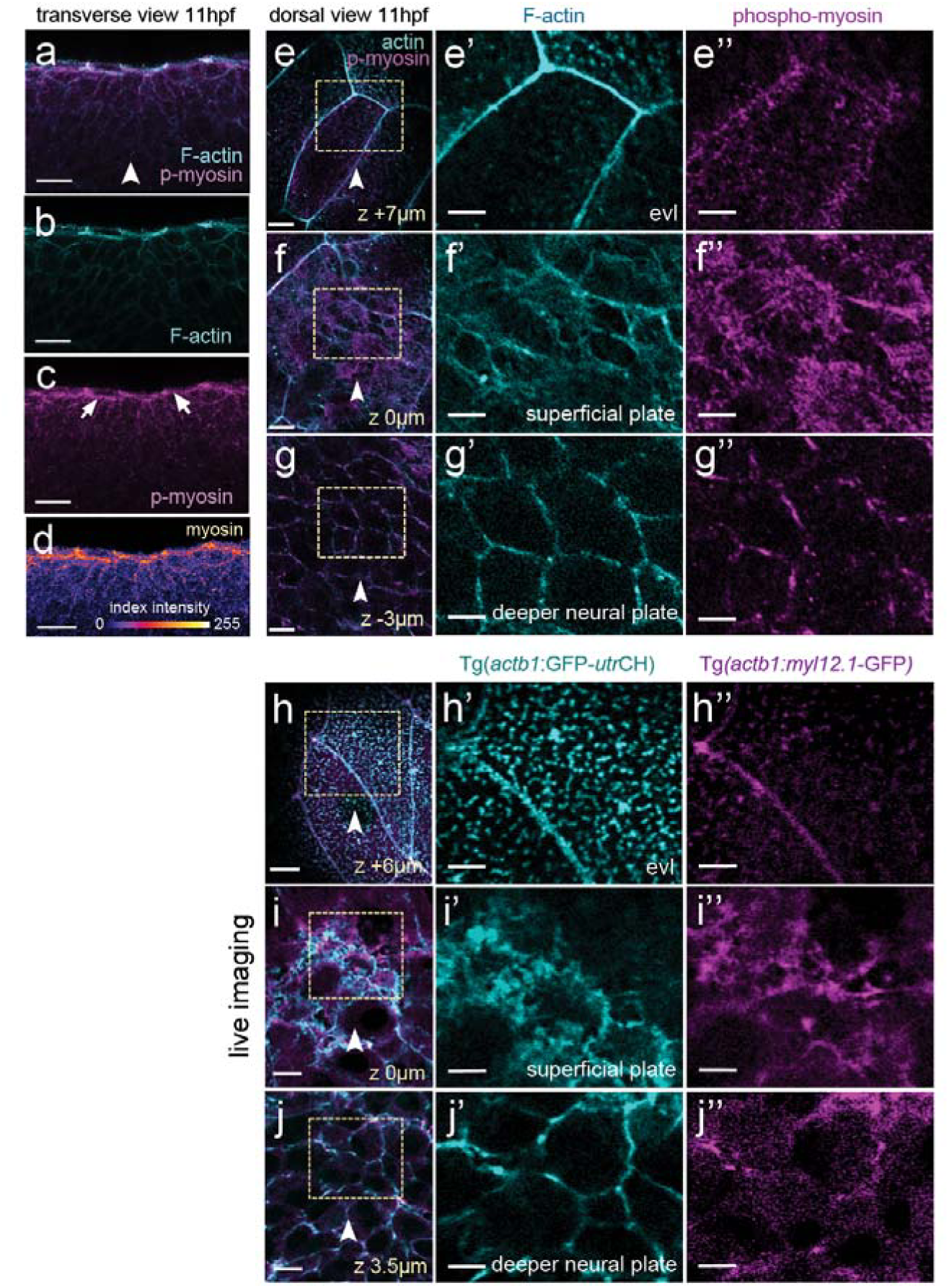
Myosin and actin in the neural plate. (**a-d**) Transverse confocal section of neural plate stained for phosho-myosin and F-actin. (**e-e’’**) Horizontal confocal section through EVL stained for phosph-myosin and F-actin. (**f-f’’**) Horizontal confocal section at level of dorsal surface of neural plate cells stained for phospho-myosin and F-actin. (**g-g’’**) Horizontal confocal section 3 μm below level of dorsal surface of neural plate cells stained for phospho-myosin and F-actin. (**h-j’’**) Myosin and actin revealed in living neural plate by Tg(*actb1:myl12.1*-GFP) and Tg(*actb1*:GFP-*utr*CH) transgenic embryos. Sections equivalent to panels (**e-g’’**). Scale bars are 5 μm.

Tranverse plane confocal movies of the Myosin-GFP and Actin-GFP embryos from 11 hpf to 14 hpf revealed their distribution during neural plate to neural keel morphogenesis. During internalisation at 11-13 hpf, non-muscle Myosin-II:GFP is strongly enriched at the dorsal surface of medial regions of neural plate (white arrows in Fig. 4a, n_embryos_ = 3, and Movie 5), and some focal accumulation can be also found at the ventral surface of the plate (cyan arrow in Fig. 4a) and scattered through the depth of the plate. As neurulation reaches the end of keel formation at 14 hpf, Myosin-II:GFP enrichment is lost from the dorsal midline position (Fig. 4a), suggesting that its activity is temporally and spatially correlated with tissue internalisation. Analysis of confocal z-stacks collected from a dorsal view confirms high levels of Myosin-II:GFP are largely restricted to dorsal surface of the neural plate cells (Fig. 4b, n_embryos_ = 3) with a marked distribution close to the superficial plasma membrane (Fig. 4b z-levels 7.5-10 μm, Fig. 4c). In contrast to Myosin-II:GFP localisation however, the distribution of Actin:GFP is somewhat more uniform across both the dorsoventral axis and mediolateral axis of the neural plate (Fig. 4e). There is a slight bias of Actin:GFP to the dorsal side of the neural plate but, in contrast to Myosin-II:GFP, the Actin:GFP gradient is much shallower within the first 40μm from the dorsal surface (Fig. 4e). To investigate Myosin-II function during internalisation we made use of the specific Myosin-II inhibitor Blebbistatin^26^. The movements of convergence of the neural plate and neural keel formation were inhibited by 50 μM Blebbistatin (Fig. 4f,g). Moreover, we found that abrogating Myosin-II activity during this developmental time, leads to a complete block of dorsal cell surface constriction that usually characterises medial neural plate cells (Fig. 4h,i, n_embryos_ = 3, n_medial cells_ = ∼35, n_lateral cells_ = ∼40). Thus Myosin-II is a significant driver of superficial cell constriction and the movements of convergence and keel formation.

**Figure 4.**
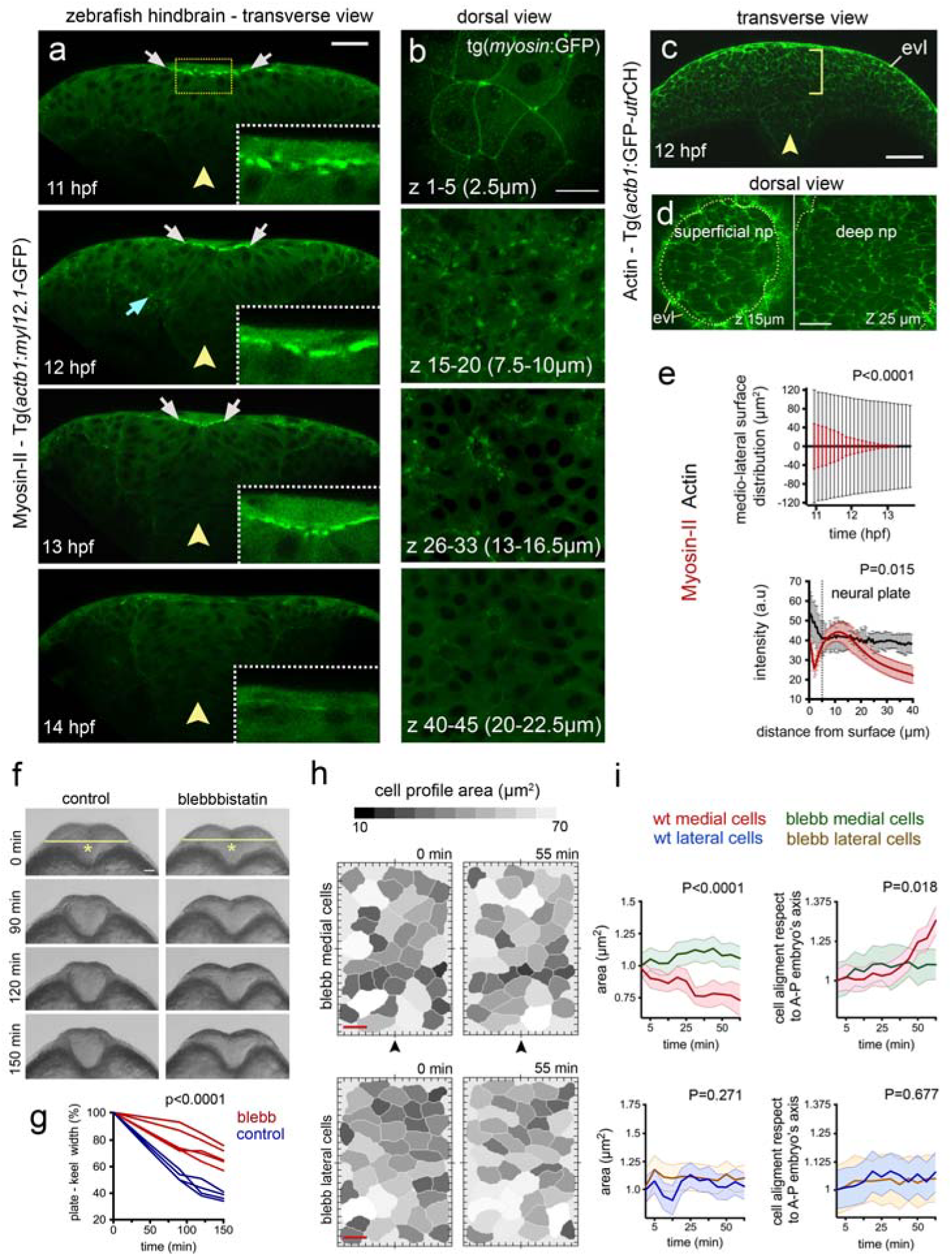
Myosin distribution and function during convergence and internalisation. (**a**)Transverse confocal images depicting the distribution of the Myosin-II:GFP fusion protein using Tg(*actb1:myl12.1*-GFP)^24^ transgenic line during zebrafish neural plate internalisation. White arrows in (a) indicate medial accumulation of Myosin-II:GFP, while cyan arrow shows basal accumulation. Insets show dorsal medial zone at higher magnification. Yellow arrowheads indicate midline position. Scale bar is 20μm. (**b**) Dorsal view confocal images of Myosin-II:GFP expression at different depths. Top image is in EVL (the superficial non-neural epithelium), then successive depths through the neural plate. Each image is a maximum intensity projection of 5 confocal slices. Scale bar, 10μm. (**c**) Representative transverse view confocal images showing actin expression by using zebrafish Tg(*actb1*:GFP-*utr*CH)^25^ reporter line during early internalisation. Arrowhead shows midline and brackets indicate superficial-deep axis for quantification in (e, bottom graph). Scale bar, 30μm and arrowhead indicates midline. (**d**) Dorsal view confocal images of Actin:GFP distribution at dorsal surface of neural plate and 10 μm deeper. Scale bar is 20μm. (**e**) Top graph: quantification of medio-lateral surface distribution of Myosin-II:GFP (red) versus Actin:GFP (black) throughout internalisation. Bottom graph: quantification of Myosin-II:GFP (red) and Actin:GFP (black) pixel intensity distribution along the superficial (EVL first 5μm) to deeper (ventral) axis of the neural plate (see methods for quantification procedure). A.U, indicates arbitrary units. (**f**) Transverse views of neural morphogenesis in control and Blebbistatin treated embryos. Scale bar is 20μm. (**g**) Quantification of neural plate to keel transition in control and Blebbistatin treated embryos. (**h**) Dorsal cell surface profiles in medial and lateral zones of neural plate in control and Blebbistatin treated embryos. Scale bar, 10μm and time is in minutes. Arrowheads indicate position of the tissue midline. (**i**) Quantification of dorsal cell surface profiles and alignment of dorsal surface areas to embryonic axis in control and Blebbistatin treated embryos (n_embryos_ = 3, n_medial cells_ = 35, n_lateral cells_ = 40). In all graphs, bars indicate SEM.

### Loss of *cdh2* leads to aberrant cell and tissue dynamics during internalisation

The generation of traction through Myosin-II dependent contractility during epithelial morphogenesis is thought to depend partially on cell-cell adhesion molecules like Cadherins^9,27^ and through adaptor proteins including α-Catenin^28,29^. Since Myosin-II drives dorsal cell constriction during neural plate internalisation and previous work shows a requirement for Cdh2 in the early stages of zebrafish neural tube morphogenesis, it is likely that the tissue and cell behaviours of internalisation should be defective in the previously identified *cdh2* mutant *parachute* (*pac*)^21,22^. Quantification of cell and tissue behaviours from transverse view imaging at prospective hindbrain levels shows that at 10-11 hpf, *cdh2/pac* mutant embryos start neurulation with a fairly normal neural plate morphology similar to wild-type animals (Fig. 5a,a’). However, by neural keel stages (12-13 hpf), *cdh2/pac* mutants display abnormal neural plate internalisation leading to a consistent and highly-reproducible aberrant T-shape or “parachute” morphology^21,22^, leading to aberrant neural tube organisation (Fig. 5a.a’, n_embryos_= 5, Fig. S1, and Movie 6). Quantification confirms *cdh2* is required to complete internalisation as *pac* mutants deepen only 62% of the neural tissue as compare to wild-type (∼wt 138μm vs ∼*cdh2/pac* 85μm, n_embryos_= 4 wt, 4 *cdh2/pac*, P◻<◻0.0001, Fig. 5b). To better define Cdh2 dependent cell behaviours we reconstructed nuclei tracks in wt and *pac* mutants. Abnormal cell behaviours were evident in *pac* neural plates from 9hpf through to 12hpf (Fig. 5c,d). The most prominent defects were seen at 11 and 12hpf when convergence of lateral cells towards the midline (Fig. 5e) and the inward trajectory of medial cells (Fig. 5f) are severely compromised. In contrast to wt nuclei that show clear differences in behaviour at lateral and medial locations, the tracks of nuclei of *pac* cells appear very similar at both lateral and medial locations (Fig. 5e,f), thus the distinctive behaviours of cells at the midline is lost. The speed of movement of medial *pac* nuclei is slower than wt nuclei, and lateral *pac* cells are initially slightly faster than wt cells but by 12hpf are significantly slower than lateral wt cells (Fig. 5g). Although loss of a cell adhesion protein might be expected to alter tissue compaction, the distance between neighbouring nuclei is not significantly different between wt and *pac* cells, showing the density of cells is unchanged (Fig. 5h).

**Figure 5.**
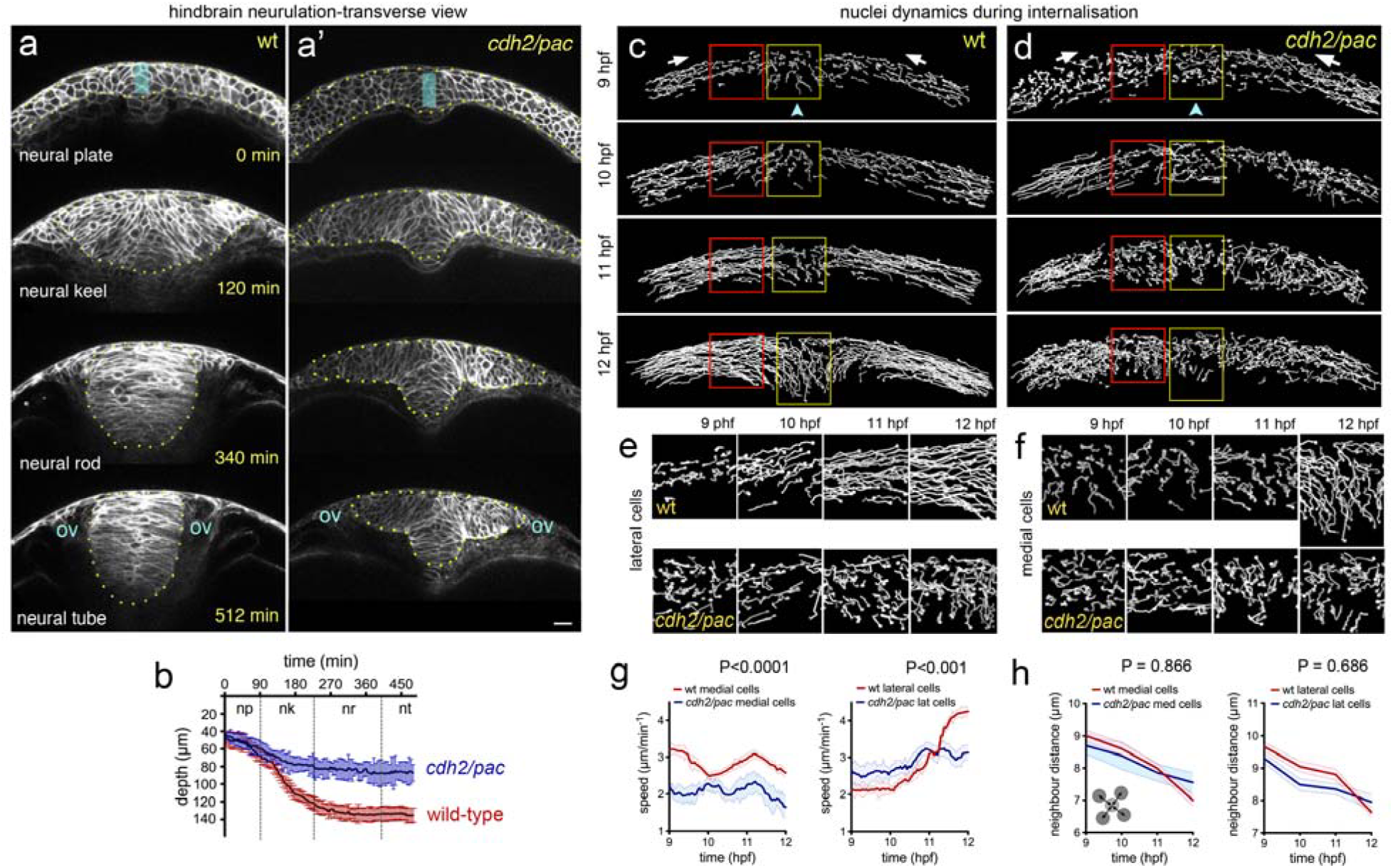
Defective cell and tissue internalisation in *cdh2/pac* mutants. (**a-a’**) Transverse confocal time-lapse images of tissue internalisation between wild-type (a) and *cdh2/pac* mutant (a’) embryos at hindbrain level. Cell membranes labelled with GFP-membrane. ov, otic vesicle. Yellow dots outline neural tissue. (**b**) Comparative internalisation dynamics between wild-type (red) and *cdh2/pac* (blue) embryos as measured by depth of the neural primordium at its midline (cyan bar in a, a’). While in wild-type embryos, internalisation is responsible to deepens up to 70% of the neural primordium (~ 42μm/~ 138μm on average), *cdh2/pac* mutants average only 62% to wt values (~ 44μm/~ 85μm, n_embryos_ = 4 *cdh2/pac*. Average depth; 85μm *cdh2/pac* vs 138μm wt). (**c** and **d**) Representative transverse images of nuclei tracks in wild-type (c) and *cdh2/pac* (d) neural plate cells during convergence and internalisation stages. *Cdh2/pac* nuclei show defective convergence and internalisation. (**e**) Higher magnification of lateral nuclei tracks taken from red boxes in (c) and (d). (**f**) Higher magnification of medial nuclei tracks taken from yellow boxes in (c) and (d). (**g** and **h**) Quantification of nuclei speeds and density between wild-type and *cdh2/pac* (n_embryos_ = 3, n_average of medial cells analysed_ = 90, n_average of lateral cells analysed_ = 80, 5 z-slices analysed in each embryo). Scale bar in (a), 20μm. Bars indicate SEM.

### Cdh2 is enriched at superficial surface during internalisation

We next asked whether Cdh2 protein localisation was consistent with a role in mediating Myosin-II dependent constriction of dorsal cell surfaces. We studied Cdh2 localisation using the BAC-reporter TgBAC(*cdh2:cdh2*-tFT), which faithfully recapitulates zebrafish Cdh2 expression^30^. The tandem fluorescent timer (tFT) technology coupled in TgBAC(*cdh2:cdh2-*tFT) comprising a rapidly maturing superfolded GFP and a slower maturing red fluorescent counterpart (tagRFP) has been used to assess protein life-time turnover, including Cdh2 maturation^30,31^. We were only able to detect robust expression of the more stable Cdh2-RFP in cell-cell junctions after internalisation at rod stages of development (14-15 hpf, Fig. S2), however Cdh2-GFP expression is readily detected from neural plate stages. Although Cdh2-GFP is present at low levels throughout the neural cell membranes, there is markedly higher expression at the dorsal surface of the neural plate (Fig. 6a,b, n_embryos_ = 4). In addition, by imaging from the dorsal surface we found Cdh2-GFP is particularly enriched at cell-cell and multi-cellular interfaces during dorsal surface constriction events close to the neural plate midline (arrowheads in Fig. 6c,c’).

**Figure 6.**
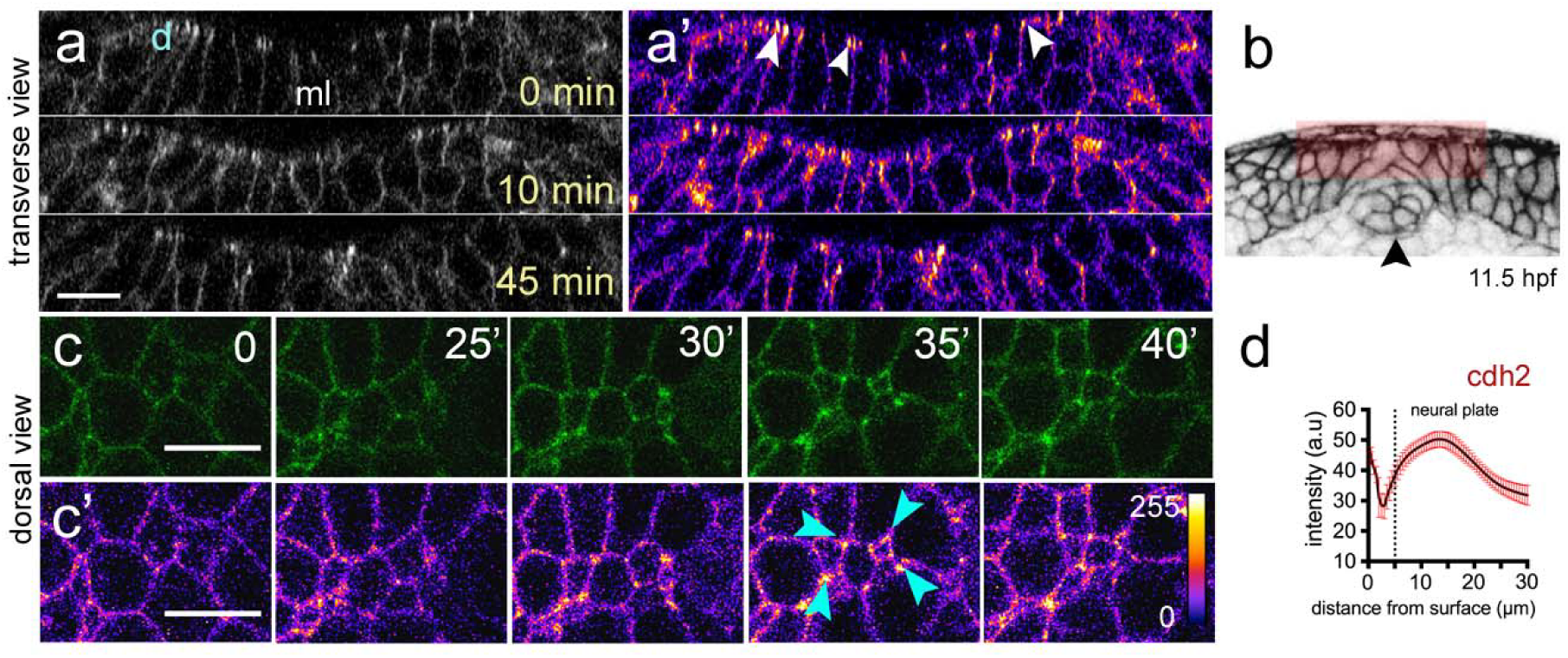
Cdh2 distribution during neural plate internalisation. (**a, a’**) Confocal transverse plane views of Cdh2-GFP at the dorsal surface of the neural plate. Cdh2-GFP is expressed throughout the cell membranes but is enriched at the dorsal surface (arrowheads). d, indicates dorsal surface. (**b**) Diagram indicating location of transverse views shown in (a, a’). (**c-c’**) Time-lapse frames of confocal images in dorsal view of the dorsal surface of the neural plate show that Cdh2-GFP is enriched at superficial surface in neural plate cells and at multi-cell interfaces. Fire lookup table indicating intensity values (0-255) in (c’). (**d**) Quantification of Cdh2-GFP intensity at different depths of the neural plate (n_embryos_ = 4). Scale bars, 10μm.

### Depletion of Cdh2 reduces dorsal contraction of medial neural plate cells

The orientation of tissue contractility and force transmission depends on the integrity of intercellular adhesion molecules to mechanically couple cells within a giving tissue^9,32^. While this phenomenon has been described in folding insect epithelia^9^, it is not entirely clear to what extent this is conserved in vertebrate neural tube morphogenesis. To explore the interrelationship of Cdh2 and Myosin-II contractility during neural plate internalisation, we studied Myosin-II:GFP dynamics in *cdh*2 deficient embryos. Reduction of Cdh2 was achieved by using a previously characterised antisense morpholino (MO), which faithfully phenocopy the *cdh2* mutant *pac* ^21^. We found that although Myosin-II:GFP is still enriched at the dorsal surface of the cdh2 MO neural plate (arrows in Fig. 7a,a’, and Movie 7) its distribution is different to wild-type embryos. Elevated Myosin-II:GFP expression is found in the medial region of the wild-type neural plate (Fig. 4a), but this preferential medial distribution is absent from *pac* embryos (Fig. 7a,b). This abnormal Myosin-II:GFP localisation is maintained into later stages of neural plate morphogenesis (Fig. 7a,b,d). To test whether reduced Cdh2 function and an abnormal Myosin-II:GFP distribution impacts the dorsal cell surface contraction events of normal internalisation we monitored dorsal cell surface profiles in Cdh2 deficient neural plates (Fig. 7e-g). In contrast to wild-type animals where internalisation is largely restricted to cells close to the midline, reduction of Cdh2 results in random cell internalisation events across the entire width of the dorsal surface of the neural plate (Fig. 7h, n_embryos_ = 5 *cdh2/pac*, n_cells_ = ∼120 per assay, and Movie 8). Individual internalisation events were confirmed by transverse view imaging (Fig. 7c). Furthermore the dorsal surface constriction of medial neural plate cells is reduced in Cdh2 depleted embryos, while constriction of lateral cells is increased (Fig. 7i,j). The orientation of the cells’ dorsal surface profile relative to the embryo’s axis is also slightly altered from wild-type (Fig. 7i,j).

**Figure 7.**
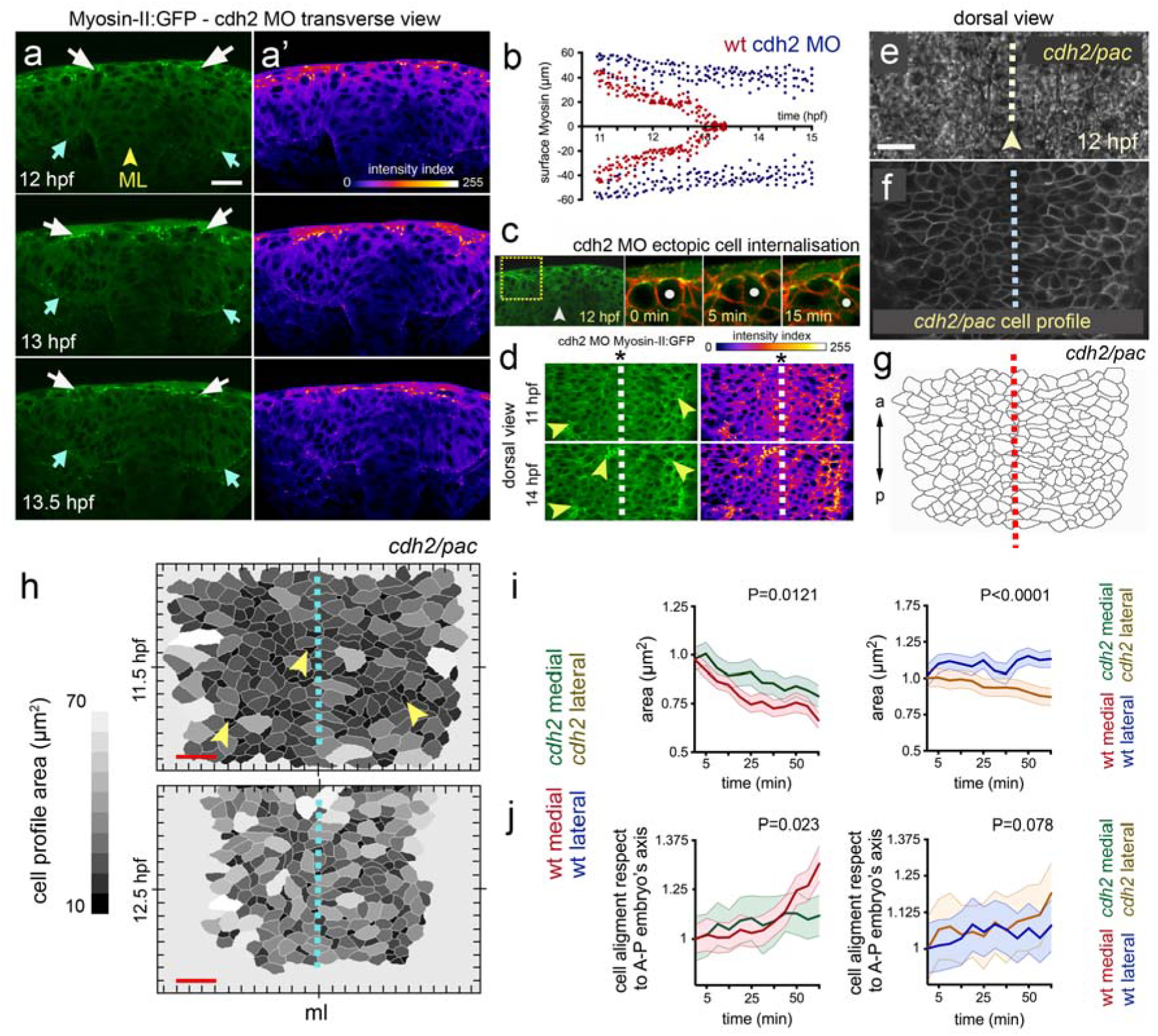
Cdh2 is required to organise midline Myosin-II during internalisation. (**a**) Frames from a transverse confocal time-lapse showing surface distribution of Myosin-II:GFP during cdh2 MO deficient neural plate internalisation (left panel). White arrows show surface Myosin-II:GFP distribution, while cyan arrows indicate basal Myosin-II:GFP accumulation. (**a’**) Fire lookup table indicating intensity values (0-255). (**b**) Comparative quantification of medio-lateral surface Myosin-II:GFP distribution between wild-type (red) and cdh2 MO (blue) during internalisation (n_embryos_ = 5 wt, 4 cdh2 MO, P<0.0001). (**c**) Representative example of ectopic cell internalisation in cdh2 MO(n_embryos_ = 3, n_ectopic cells internalisation_ = 28). (**d**) Left panels: single dorsal confocal frames showing abnormal midline Myosin-II:GFP distribution in cdh2 MO embryos (arrowheads). Right panels: fire lookup table, showing intensity profile. Dashed line represent midline. (**e-g**) Dorsal picture of cdh2 MO neural plate. (**e**) Bright field dorsal view images in *cdh2/pac* tissue by 12hpf. Scale bar is 20μm. (**f**) Single dorsal confocal slice depicting cells’ dorsal area profiles in *cdh2/pac* neural plate labelled with membrane-GFP (gray). (**g**) Automated cell area profile reconstruction from (f). a = anterior, p = posterior. Arrowhead and dashed lines in (e),(f) and (g) represent tissue midline. (**h**) *Cdh2/pac* mutant dorsal cell surface profiles during defective internalisation. Arrowheads indicate cell constriction events and dashed line represent tissue midline (ml). Scale bar, 10μm and time is in hpf. (**i-j**) Quantification of dorsal cell area profiles during *cdh2/pac* mutant internalisation. (**i**) Mean cell surface area between medial (wt red, *cdh2/pac* green left, P=0.0121), and lateral cells (wt blue, *cdh2/pac* orange, P<0.0001). (**j**) Comparative cell aligment along the anterior-posterior embryo’s axis between medial (wt red, *cdh2/pac* green left, P=0.023), and lateral cells (wt blue, *cdh2/pac* orange, P=0.078).

## Discussion

Zebrafish embryos generate a neural tube by a complex array of cell behaviors^10,11,12,15^. Initially this involves the movement of a large body of cells towards the dorsal midline^14,15,16^. Then tissue internalisation at the midline transforms the neural plate into a neural keel and then solid cylindrical neural rod structure. These highly organised collective movements of dorsal midline cells are dependent on extrinsic cues from mesoderm and extracellular matrix^15,16,17^, as well as cell-cell interactions between the neural cells themselves^13,14, 17, 21, 22^. While tissue internalisation is a fundamental step in this morphogenetic process, its cellular and molecular nature has remained rather elusive. In this work we have shed light on some of the cellular events of neural plate cell internalisation in the zebrafish. This step is driven by previously uncharacterised cell surface constriction events in the medial zone of the neural plate. We show these superficial constriction events are dependent on Cdh2 and Myosin-II.

Unlike other vertebrates the teleost neural plate is not a columnar epithelium and does not transform into a neural tube by the conventional primary neurulation mechanism of epithelial folding. Additionally the distribution of polarity proteins within the teleost neural plate^18^ and its unusual cell behaviours have led to the view that it is not an obviously epithelialised tissue. Despite this, our study suggests internalising zebrafish neural plate cells deploy a similar strategy and molecular machinery to that widely used in epithelial tissue infolding events. Central to this in other systems is the presence of a contractile activity of non-muscle Myosin-II and cell-cell adhesion machinery of Cadherins at cell and tissue surfaces^4,7,9,19,20^. During epithelial morphogenesis in flies, cortical Myosin-II has been shown to exhibit a distinctive pulsatile behaviour in internalising cells while assuming a more interconnected network at tissue level^7,9^. Our live imaging analyses reveal that during zebrafish neural plate internalisation, Myosin-II shows a transient enrichment at the superficial surface of medial neural plate cells. While at the moment we do not know the higher resolution organisation and dynamics of Myosin-II activity in individual cells of the fish neural plate, we hypothesise that Myosin-II activity exerts a mechanical tension across the dorsal surface of medial neural plate cells to progressively constrict their superficial surface and mediate cell internalisation. We find cell internalisation appears to occur on a cell by cell basis rather than a coordinated internalisation of neighbouring cells.

Myosin-II dependent forces at the cell cortex are thought to be organised and transmitted through the tissue by cell-cell surface molecules including Cadherin family^9,27,29^. During Drosophila mesoderm invagination, precise accumulation of Myosin-II at the ventral midline surface greatly depends on adherens junctions^36^. Disruption of these cell-cell adhesion molecules during insect gastrulation, result in broader Myosin-II expression at the midline surface and impaired tissue invagination^36^, which is reminiscent to our observation of Myosin-II distribution in neural plate depleted of Cdh2. Similar to E-cadherin function^9^, lack of N-cadherin during insect epithelial morphogenesis result in abnormal tissue remodeling, impaired Myosin-II organisation and defective cell anisotropy^37^. During neurulation in mice and frogs, the depletion of Cdh2 results in defective Myosin-II expression and aberrant neural tube development^34,35,38^. Although the teleost neural plate is structurally different to other vertebrates the impaired convergence and internalisation of neural cells in the zebrafish *cdh2/pac* mutant has some similarities to an open neural tube phenotype. Here we show Myosin II contractility is a significant driver of cell surface constriction at the dorsal surface of neural plate and cell surface constriction is defective in Cdh2 mutants. Thus Cdh2 is required to mediate the dorsal cell surface remodeling required in the medial zone of the neural plate to effect cell internalisation.

Our current work has focused on the cell constriction behaviours at the dorsal surface of the zebrafish neural plate and the contribution of deeper cell behaviours to neural plate morphogenesis remain unknown. In other systems apical constriction is often accompanied by expansion of basal cell surface resulting in wedge-shaped cell morphologies and this is thought to be a paradigm for tissue folding in a wide range of developmental contexts^3,4,6^, including vertebrate neural tube formation^4,19,20^. Our preliminary analysis of cell morphologies from ubiquitously labelled cells (Fig. 1) suggests zebrafish cells acquire spindle shaped morphologies during internalisation rather than wedge-shaped morphologies, and suggest most cells do not stretch from the dorsal surface to the ventral surface of the plate during this behavior. It seems likely that cell elongation in the medial zone of the neural plate contributes to deepening of the tissue as it transforms into the neural keel, but an accurate description of individual cell morphologies and behaviours awaits more comprehensive analysis of mosaically labelled tissues. It will also be important to understand the nature of the junctions between cells at the dorsal surface of the plate and this will probably require a 3D-EM characterisation^33^.

In conclusion, although its organisation and cell behaviours^10,11,12,14,15,16,17,18,22^ mean the epithelial nature of the teleost neural plate remains somewhat enigmatic, our results suggest it none the less employs cell surface constriction strategies, based on Cadherin adhesions and Myosin II contractility, typically used by epithelia for internalisation events. Moreover, we hypothesise that this epithelia-like remodeling has to be transient in the fish neural plate in order to allow the subsequent repertoire of cell interdigitation and mirror-symmetric divisions that occur across the neural midline during keel and rod stages^12,13,17,22^.

## Methods

### Ethics statement

All experimental procedures involving animals were approved by the College Research Ethics Committee at King’s College London (London, UK) and covered by the Home Office Animals (Scientific Procedures) Act 1986 (ASPA) project licence, and by the Comité de Etica y Bioética DID-UACh (Bioética en uso de Animales en la Investigación, DID-UACh). All methods were performed in accordance with the relevant guidelines and regulations.

### Zebrafish husbandry and strains

Wild-type, transgenic and mutant adult zebrafish were maintained under standard conditions^39^ on a 14-hour photoperiod at the KCL, and UACh Fish Facilities, respectively. Embryos were collected from timed matings and raised at 28.5°C in fish water or E3 embryo medium^39^. Animals were staged according to published morphological criteria^40^ and stages are given in terms of hours post fertilisation (hpf). The following strains were used in this study: wild-type (wt) TL and AB, *cdh2*^fr7(21)^, Tg(*actb1:myl12.1*-GFP)^24^, Tg(*actb1*:GFP-*utr*CH)^25^, and TgBAC(*cdh2:cdh2-*tFT)^30^.

### mRNA synthesis and microinjections

PCS2+ expression vectors expression carrying membrane-localised green fluorescent protein GFP (mGFP), membrane-localised red fluorescent protein RFP (mRFP), nuclei histone 2B GFP (H2B-GFP) or nuclei histone 2B RFP (H2B-RFP) constructs were linearised using restriction enzymes for 3 hours at 37ºC and precipitated at −20ºC in 70% ethanol. DNA was then washed and resuspended in RNAase free water. Sense capped mRNA was transcribed using the mMESSAGE SP6/T7/T3 Kit (Ambion) and purified later using columns (Roche). Finally, mRNA concentration was measured in nanodrop (Thermoscientific). All microinjections were performed in a dissecting microscope using a glass slide and petri dish. Injections were made at one-cell stage embryo and mRNA constructs (150 pg per embryo) were delivered using a glass micropipette with filament (Harvard Apparatus) mounted on a micromanipulator and attached to a Picospritzer^®^ (General Valve Corporation).

### Antisense morpholino oligonucleotide injections

To inhibit Cdh2 function we used a previously characterised cdh2 morpholino (MO) oligonucleotide (cdh2 MO)^21^ against (5-TCTGTATAAAGAAACCGATAGAGTT-’3), corresponding to –40 to –16 of cdh2 cDNA^21^. cdh2 MO was purchased from Gene Tools (Philomath, OR), dissolved in water to a stock solution of 1mM, and injected at 1-cell stage embryo at a final concentration of 50μM.

### Antibody staining

Whole-mount and tissue section immunohistochemistry was performed as previously described^12,16^. In this study we used mouse anti-ZO-1 (339111; Zymed) at 1:300, anti-rabbit Phospho-Myosin Light Chain 2 (Ser19) (3671; Cell Signaling) at 1:20 in 2.5% normal goat serum (Sigma). For secondary antibodies we used anti-mouse Alexa 488 and 633 (Molecular Probes) at 1:1,000 in 2.5% normal goat serum. For F-actin labelling we used Phalloidin (A12379; Invitrogen) at 1:1,000 in 2.5% normal goat serum. For analysis of neural tube organsition tissue sections were made every 14 mm on a HM 560 MV Micron microtome.

### Blebbistatin treatment

To block Myosin-II activity, wild-type or Tg(*actb1:myl12.1*-GFP) embryos at 10 hpf were put in a small dish containing either 50 μM or 100 μM of Blebbistatin dissolved in E3 medium and raised at 28.5°C. Transient Blebbistatin treatments were kept until 12 hpf while embryos were imaged by confocal microscopy. After pharmacological treatments embryos were washout from Blebbistatin and their morphology was assessed by 24 hpf.

### Time-lapse imaging

For supplementary movie 2 and data figure 4, we used Bioemergence/Mov-IT workflow analysis^23^. Dechorionated wild-type and *cdh2/pac* embryos were transferred to 0.5% low melting point agarose (Sigma) and mounted dorsally at level of hindbrain in a 3 cm petri dish with glass containing a 0.5 x 0.78 mm^2^ hole at the centre and surrounded by teflon mold. Image acquisition was performed at 28.5°C on an upright Leica SP5 confocal microscope using an Olympus objective 20X/0.95 NA water dipping lens. For imaging field size, we recorded at 351×351 μm for XY and 171μm of Z depth, with 0.69×0.69×1.05μm^3^ voxel size and time step of 2.5m. After imaging recording, embryos were assessed for general morphology and potential neural tube defects. For all other movies, dechorioanted embryos were mounted in 0.8% low melting point agarose (Sigma) in embryo medium (E3). Transgenic animals and embryos expressing RFP/GFP-tagged membrane or nuclei constructs were imaged in either transverse or dorsal view at the level of the prospective hindbrain using the otic ear as anatomical landmark at 18 hpf. For transverse view registration images were taken 3-5 μm apart while for dorsal recording confocal images were taken 0.25 μm apart. In both cases, time-lapses were acquired on an upright Leica TCS SP5 LSM using a Leica 25x 0.95 NA water dipping lens and a temperature chamber at 28.5°C. Z-stacks were collected at 2-4-minute intervals, starting at 10 hpf and continuing in some cases up to 20 hpf.

## Imaging processing

### Center detection and automatic tracking at Bioemergence

Collections of .vtk files were uploaded to Bioemergences workflow^23^. Nucleus centre detection and cell tracking was performed using Mov-IT^23^. Automated 4D tracking of approximately 3500 cells within a 351×351 μm volume was performed^16^. For reconstruction of transverse movies, groups of ~100 neural plate cells were manually selected in standardised ROI in the developing hindbrain near the otic vesicle. Automated nuclei tracking was manually validated.

### Cell dynamics

To quantify cell shape changes during internalisation, we used FIJI imaging software^41^ and customised Matlab code. Briefly, segmented cells were automatically obtained with FIJI by using the “trainable weka segmentation”^42^ plugin. Segmented binary images were then used in our customised Matlab code (MathWorks, Cambridge, UK) to extract cell parameters using major/minor axis on each cell computed using second order image moments (*regionprops* function in Matlab). A gray scale was used to quantify cell area (black = minimum cell area, white = maximum cell area, for each movie). To quantify cell aspect ratio independently of area, we divided the major axis by the minor axis to build an eccentricity index. For cell anisotropy measurements, we computed the absolute value of the relative angle of the major cell axis with respect to the anterior-posterior embryo axis, where the later was obtained manually from bright-field images.

### Pixel intensity analysis

To quantify pixel intensity for Figures 4 and 6, the following approach was used: i) a consistent imaging set up was kept in all experimental conditions (stage of development, day of embryo imaging, construct concentration, confocal settings, laser power, offset, pinhole and averaging), ii) a histogram of all pixel intensities (256 gray levels) was calculated using three small region of interest (ROI, 3μmx3μm) in 20 cells at each individual z-level. For dorsal surface measurements, we calculated “true signal” by averaging the intensity from maximum intensity projections of 3-4 representative z-slices, including EVL signals when necessary (i.e. Myosin-II:GFP). Background pixel intensity (avoiding membrane signal) was subtracted from the average values. Images were analysed using FIJI software^41^ and presenting intensity in Fire LUT colour code when necessary (values, 0-255).

### Statistics

To test for significance between mean values, student’s t-test p values were calculated using non-parametric Mann-Whitney post-test (Matlab, Cambridge, UK statistic toolbox and Graph Pad, La Jolla, CA, USA). We used a probability of 0.05 for significance. All the error bars shown in figures are SEM (standard error of the mean). For each experiment, we normally used 3-6 embryos. Graphical representations were performed using GraphPad Prism programme.

## Supplementary Figure Legends

**Figure S1.**
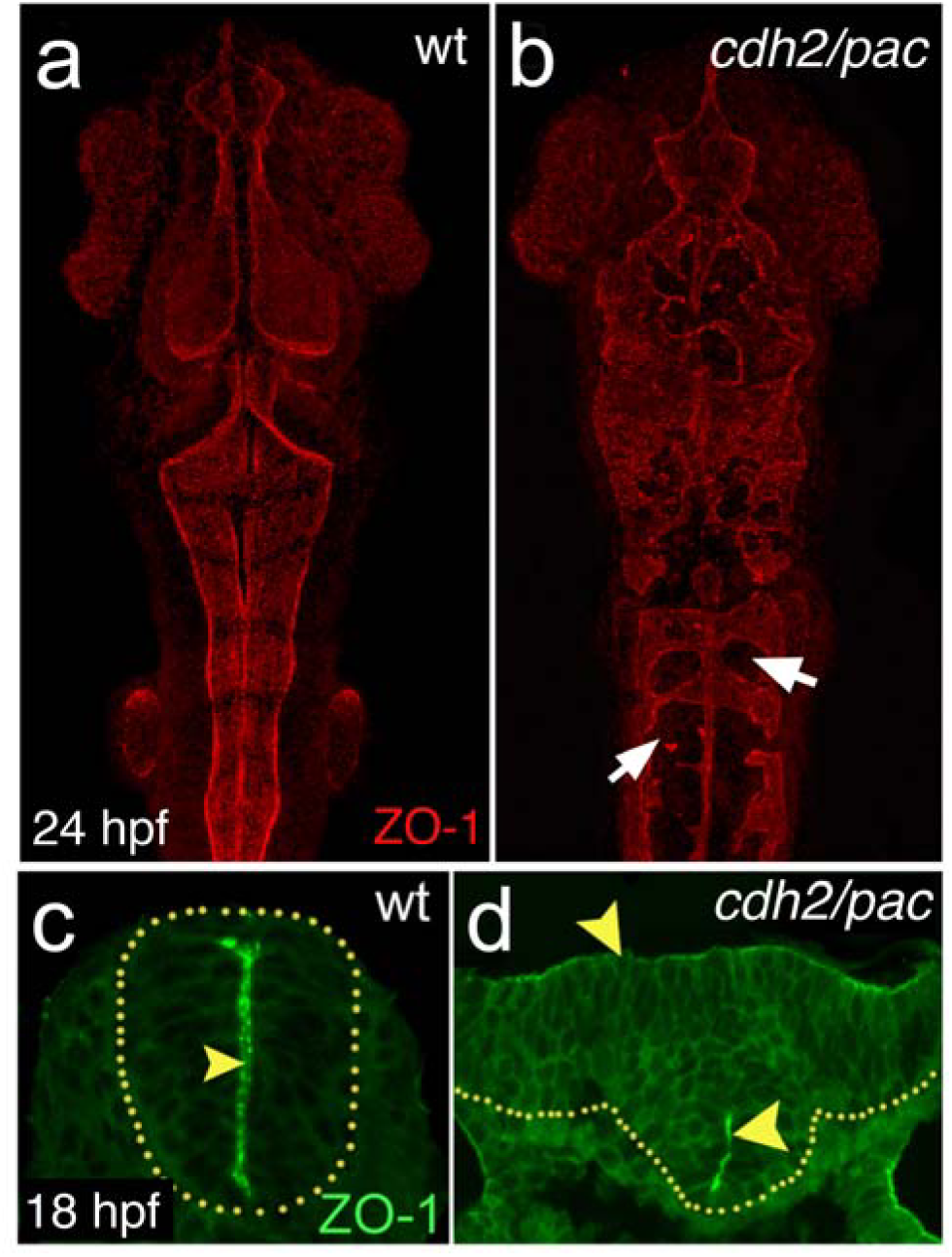
Depletion of *cdh2* leads to defective neurulation and abnormal apico-basal polarity. (**a-b**) Maximum projections of dorsal confocal pictures showing abnormal neural tube morphology and disorganised apical midline seam specialisation in *cdh2/pac* by 24 hpf, as revealed by apical marker, ZO-1 (arrows). Anterior is up. (**c-d**) Cross-section analysis in *cdh2/pac* mutants reveal disrupted apical midline seam organisation (arrowheads) at hindbrain levels.

**Figure S2.**
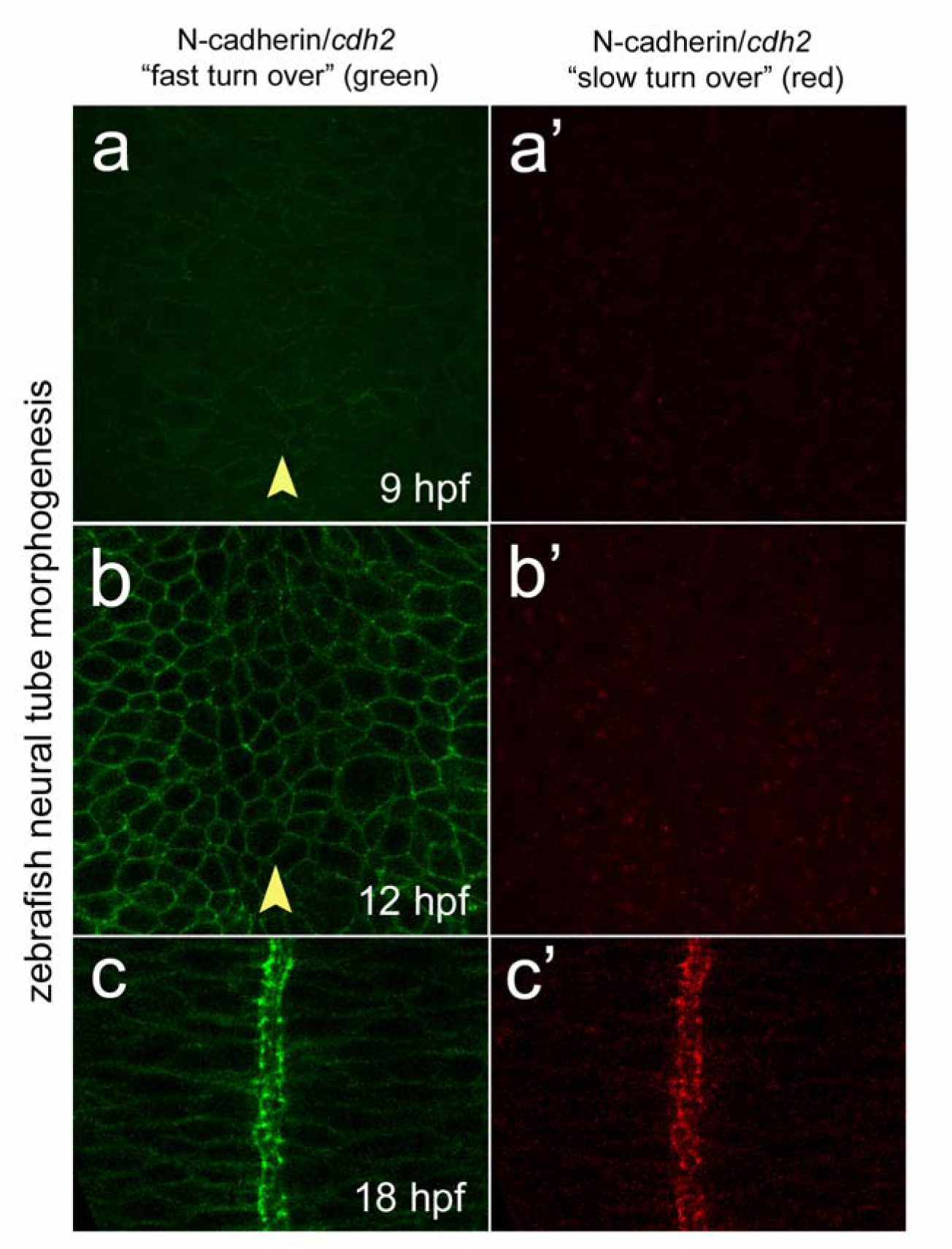
Cdh2 turnover through zebrafish neurulation. (**a**) Dorsal single confocal picture of zebrafish neural plate at 9 hpf showing Cdh2-fast/dynamic “green” turnover in TgBAC(*cdh2:cdh2-*tFT)^24^. (**a’**) Dorsal single confocal picture of zebrafish neural plate at 9 hpf showing n-cadherin slow/stable “red” turnover in TgBAC(*cdh2:cdh2-*tFT). (**b**) Dorsal confocal picture of zebrafish neural plate at 12 hpf showing Cdh2 “green” turnover. (**b’**) Dorsal confocal picture of zebrafish neural plate at 12 hpf showing Cdh2 “red” turnover. (**c**) Dorsal confocal picture of zebrafish neural tube at 18 hpf showing Cdh2 “green” turnover. (**c’**) Dorsal confocal picture of zebrafish neural tube at 18 hpf showing Cdh2 “red” turnover. Arrowhead indicates tissue midline.

## Movie Legends

**Movie 1. Tissue morphogenesis during zebrafish hindbrain neurulation.** Cells expressing GFP-caax to label membranes. Movie collected from 10-17.5 hpf and covers neural plate convergence and internalisation, transition to neural keel and then to neural rod. Arrowhead indicates midline and ov indicates otic vesicles as anatomical landmark for hindbrain. Dorsal to top. Frames are every 5 minutes. Time is in hours and minutes from start of movie.

**Movie 2. Nuclei tracking reconstructed from embryo expressing H2B-GFP. during zebrafish neural plate internalisation.** Dorsal view movie of a wild-type embryo from approximately 10-14 hpf covers convergence and internalisation of neural plate. Anterior to top. Arrowhead indicates midline. Frames are every 2.5 minutes.

**Movie 3. Dorsal cell surface dynamics in medial zone of neural plate.** Segmented representation of the dorsal surface profile of medial neural plate cells derived from membrane labelled cells, individual profiles cannot always be followed from frame to frame. Movie from 11-12 hpf covering period of cell internalisation in medial zone. Colour coded so larger cell surfaces are light grey and smaller surfaces are darker greys. Arrowhead indicates midline. Frames are every 5 minutes.

**Movie 4. Dorsal cell surface dynamics in lateral zone of neural plate.** Segmented representation of the dorsal surface profile of lateral neural plate cells derived from membrane labelled cells, individual profiles cannot always be followed from frame to frame. Movie from 11-12 hpf. Colour coded so larger cell surfaces are light grey and smaller surfaces are darker greys. Arrowhead indicates direction of movement towards medial zone. Frames are every 5 minutes.

**Movie 5. Myosin-II localisation during plate to keel transition.** Single transverse view time-lapse movie of a transgenic Tg(*actb1:myl12.1*-GFP) zebrafish embryo. Yellow arrows indicate high levels of Myosin-GFP at the dorsal surface of medial zone during period of internalisation. Arrowhead indicates midline. Frames are every 5 minutes.

**Movie 6. Tissue morphogenesis during neural plate morphogenesis in *cdh2/pac* mutant embryo.** Transverse view time-lapse movie from 10-15.15 hpf. Notice that during neural keel stages tissue fails to internalise and the neural anlage assumes a “T-shape” configuration. Arrowhead indicates midline and ov indicates otic vesicles as anatomical landmark for hindbrain. Frames are every 5 minutes. Time is in hours and minutes from start of movie.

**Movie 7. Abnormal Myosin-II localisation during neural plate morphogenesis in Cdh2 deficient embryo.** Transverse view time-lapse movie of a transgenic Tg(*actb1:myl12.1*-GFP) zebrafish embryo injected with cdh2 morpholino. Yellow arrows indicate high levels of Myosin-GFP across a wide expanse of dorsal surface of neural plate. Arrowhead indicates midline. Frames are every 5 minutes.

**Movie 8. Dorsal cell surface dynamics across neural plate of *cdh2/pac* mutant embryo.** Segmented representation of the dorsal surface profile of neural plate cells derived from membrane labelled cells, individual profiles cannot always be followed from frame to frame. Movie from 11.5-12.5 hpf. Colour coded so larger cell surfaces are light grey and smaller surfaces are darker greys. In contrast to wt embryos, small cell surface area profiles are more evenly spread across mediolateral axis. Arrowhead indicates midline. Frames are every 5 minutes.

## Acknowledgements

We thank members of Araya’s lab for helpful discussion and comments on the manuscript. This work was supported by grants from Dirección de Investigación y Desarrollo UACh S-201613 to CA and LC, FONDECYT 11161033; FONDEQUIP EQM140119; ACT1402; P09-015-F; CORFO (16CTTS-66390); DAAD (57220037, 57168868) to MC, CONICYT/ECOS-SUD C13B03 to CA, TS and NP, and Wellcome Trust to JC.

## Author Contributions

C.A. conceived and designed the study. C.A., and J.C. analysed all experimental results. H.H. helped to analyse data from figures 1 and 5 and generated time-lapses in figures 3 and 5. M.C. implemented algorithm to analyse figures 2, 3 and 7. L.C. helped to analyse figure 1, 2 and 7. T.S. generated Bioemergence and MOVIT platforms and helped to analyse data from figure 1. C.R. helped to generate data for figure 3 and analyse data for figure 3. N.P. helped to generate time-lapse data from figures 1 and 5 and supervised Bioemergence/MOVIT analysis. C.A., and H.H. carried out the experiments. C.A., and J.C. wrote and revised the manuscript. All authors reviewed the manuscript.

